# Polygenic risk of psychiatric disorders exhibits cross-trait associations in electronic health record data

**DOI:** 10.1101/858027

**Authors:** Rachel L. Kember, Alison K. Merikangas, Shefali S. Verma, Anurag Verma, Renae Judy, Regeneron Genetics Center, Scott M. Damrauer, Marylyn D. Ritchie, Daniel J. Rader, Maja Bućan

## Abstract

**Objective:** Prediction of disease risk is a key component of precision medicine. Common, complex traits such as psychiatric disorders have a complex polygenic architecture making the identification of a single risk predictor difficult. Polygenic risk scores (PRS) denoting the sum of an individual’s genetic liability for a disorder are a promising biomarker for psychiatric disorders, but require evaluation in a clinical setting.

**Methods:** We develop PRS for six psychiatric disorders (schizophrenia, bipolar disorder, major depressive disorder, cross disorder, attention-deficit/hyperactivity disorder, anorexia nervosa) and 17 non-psychiatric traits in over 10,000 individuals from the Penn Medicine Biobank with accompanying electronic health records. We perform phenome-wide association analyses to test their association across disease categories.

**Results:** Four of the six psychiatric PRS were associated with their primary phenotypes (odds ratios between 1.2-1.6). Individuals in the highest quintile of risk had between 1.4-2.9 times higher odds of the disorder than the remaining 80% of individuals. Cross-trait associations were identified both within the psychiatric domain and across trait domains. PRS for coronary artery disease and years of education were significantly associated with psychiatric disorders, largely driven by an association with tobacco use disorder.

**Conclusions:** We demonstrate that the genetic architecture of common psychiatric disorders identified in a clinical setting confirms that which has been derived from large consortia. Even though the risk associated is low in this context, these results suggest that as identification of genetic markers proceeds, PRS is a promising approach for prediction of psychiatric disorders and associated conditions in clinical registries.

## Introduction

Psychiatric disorders are a severe global health problem, with 7.4% of all disability adjusted life years worldwide attributable to mental or behavioral health issues(1). In the United States, 18.9% of adults have a prevalent psychiatric disorder(2), and half of all US adults are estimated to experience a psychiatric disorder in their lifetime(3). Genetic variation contributing to psychiatric disorders has been identified by multiple genome-wide association studies (GWAS) in large samples(4–8); however, due to their highly polygenic nature, no single variant or biomarker has been identified that reliably predicts the development of a disorder.

Polygenic risk scores (PRS) denoting an individual’s genetic liability for a disorder are an emerging approach that offers the possibility of acting as a biomarker for disease(9). PRS aggregate the risk carried by multiple genetic variants, weighted by the number of alleles at each contributing polymorphic locus and the effect size of those variants in a GWAS for that disorder(10). PRS have been shown to identify individuals at increased risk for non-psychiatric disorders to an extent equivalent to large-effect monogenic mutations(11), with individuals in the highest PRS categories having as much as a 25-fold increase in risk(12). In addition to their primary association with the disorder from which they were derived(13,14), PRS have been used to investigate how genetic liability for a disorder is associated with intermediate phenotypes or behavioral outcomes(15,16). However, nearly all psychiatric PRS have been evaluated in ascertained research cohorts, rather than tested in a clinical population. An exception to this was the finding by Zheutlin et al. (2019)(17) that a PRS for schizophrenia was robustly associated with a schizophrenia diagnosis in a health care setting, along with other medical diagnoses. The utility of other psychiatric PRS for risk prediction in clinical settings is unknown.

PRS also offer the potential to investigate the genetic overlap between disorders, by testing the association of PRS across multiple phenotypes. Psychiatric disorders share both symptomology and underlying genetic architecture(6,18,19). Family members of individuals with a psychiatric disorder have an increased risk for a range of psychiatric disorders not limited to the specific disorder carried by their relative(20). Individuals with psychiatric disorders also have increased rates of other medical disorders(21), although the extent to which this is driven by shared genetic etiology is unclear. Assessing whether genetic liability for psychiatric disorders is disorder-specific or has broader health implications would increase our understanding of the shared genetic etiology among different disorders.

Electronic health records (EHR) provide a wealth of phenotypic information, extending across trait domains to capture a wide picture of disease burden for each individual. Linking EHR information with genotypic data allows us to investigate cross-phenotype associations of genetic risk using a phenome-wide association study (PheWAS)(22). The use of individual-level data also permits us to explore whether the cross-phenotype associations are driven by horizontal pleiotropy, in which genetic variants convey risk independently to two different phenotypes; or vertical pleiotropy, in which genetic variants convey risk to one phenotype, which in turn raises risk for a secondary phenotype(17).

To quantify the predictive value of psychiatric PRS in an EHR setting, we generated risk scores for six common psychiatric disorders (schizophrenia, bipolar disorder, major depressive disorder, cross disorder, attention-deficit/hyperactivity disorder, and anorexia nervosa) and evaluated their association with their primary phenotype in an academic biobank - the Penn Medicine Biobank (PMBB). To identify cross-trait associations and phenotypic overlap between genetic liability for psychiatric disorders, we conducted a primary PheWAS. We performed a secondary PheWAS in which we co-varied for the primary phenotype to test whether the cross-phenotypic associations remained after adjustment for the primary phenotype. Finally, we explored whether genetic risk for other common, complex traits is associated with psychiatric phenotypes in the EHR.

## Subjects and Methods

### Penn Medicine Biobank Cohort

The Penn Medicine Biobank (PMBB) recruits participants through the University of Pennsylvania Health System by enrolling them at the time of a medical appointment. Patients give informed consent to participate by donating either blood or a tissue sample and allowing researchers access to their electronic health information. The study was approved by the Institutional Review Board of the University of Pennsylvania. As of November 2018, PMBB comprises 52,853 consented individuals, of whom 19,515 have been genotyped to date.

### Genotyping and Quality Control

DNA extracted from the blood of 19,515 total samples were genotyped in three batches: 1) 10,867 samples on the Illumina InfiniumOmniExpress-24v1-2_A1 chip at the Regeneron Genetics Center (number of single nucleotide polymorphisms (SNPs) =713,599); 2) 5,676 samples on the Global Screening Array (GSA) V1 chip (number of SNPs = 700,078) and 3) 2,972 samples on the GSA V2 chip (number of SNPs = 759,993) at the Center for Applied Genomics at the Children’s Hospital of Philadelphia. Each batch was quality controlled (QC) separately prior to imputation. Samples were removed if inferred sex from genotypes did not match their reported sex, or if they have a genotyping call rate <90%. SNPs were removed if call rate <95%, if they were palindromic, or if the SNPs were not present in the reference panel used for imputation. Following QC, from each batch we retained: 1) 10,506 individuals and 651,366 variants; 2) 5,660 individuals and 666,032 variants 3) 2,965 individuals and 700,984 variants.

### Phasing and Imputation

Genotypes for each of the three PMBB datasets were phased (Eagle v2.3) and imputed to the 1000 Genomes reference panel (1000G Phase3 v5) using the Michigan Imputation Server(23). Following imputation, the datasets were merged, matching each position based on alleles. Genotyped variants removed during imputation were manually merged back into the final dataset. In the final merged dataset, the average quality of imputation was R^2^=0.75. Genetic ancestry was calculated from common, high-quality SNPs (MAF > 0.05, missingness < 0.1) using the smartpca(24) module of the EIGENSOFT package. The datasets were split by genetic ancestry and QC steps were subsequently conducted independently for each population. As the calculation of PRS requires summary statistics from GWAS performed in a sample of the same ancestry as the target dataset, we retained individuals of European ancestry only for this analysis (N=11,524). We removed SNPs with imputation marker R^2^ < 0.7 and minor allele frequencies <0.01 to retain high quality, common SNPs. Related individuals were identified using a graph-based algorithm after applying a Pi-HAT threshold of 0.25 (to identify first or second degree relatives), and the sample that was most closely related to multiple other samples was removed (n=1,088). Individuals were removed if genotyped sex did not match their reported gender (n=85). Following QC, we retained 10,351 individuals. We used PLINK v1.9(25) to calculate ancestry specific principal components (PCs) to use as covariates.

### Polygenic risk scores

PRS were generated using LDPred (v1.0)(26) for six psychiatric disorders and 17 other traits (see Supplemental Table 1). LDPred adjusts SNP effect sizes identified in GWAS for the effects of linkage disequilibrium (LD). To reduce the computational cost of this method, we first extracted SNPs present in the HapMap reference panel(27) from the QC’d PMBB dataset (N=1,320,405). LD information from the PMBB dataset was used to generate the posterior mean effect of each SNP, conditioning on a prior for genetic architecture. The prior has two parameters – the heritability of the phenotype estimated from the GWAS summary statistics, and the estimated fraction of causal markers. Due to the highly polygenic nature of the complex disorders selected for PRS, we set the parameter representing the fraction of causal variants *ρ*=1. PRS were calculated using PLINK v1.9(25) by summing all variants, weighted by the effect size of the variant following LDPred adjustment.

### Phenotypes

We extracted International Classification of Diseases (ICD) ninth revision (ICD-9) and tenth revision (ICD-10) data for 52,853 individuals from the EHR, which consisted of 11.8 million records. We filtered on encounter type to identify records representing encounters with a physician (see Supplemental Table 2 for encounters selected). We retained data for 52,354 individuals, for 9.2 million records, with 5.3 million records being ICD-9 based, and 3.9 million records being ICD-10 based. ICD-9 codes were aggregated to phecodes using the phecode ICD-9 map 1.2(28); ICD-10 codes were aggregated to phecodes using the phecode ICD-10-CM map 1.2 (beta)(29). Individuals were considered cases for the phenotype if they had at least 2 instances of the phecode on unique dates, controls if they have no instance of the phecode, and ‘other/missing’ if they had one instance or a related phecode. In total, 1,859 phecodes were created. The final dataset included 10,182 European individuals with complete genotype, phenotype, and covariate data.

### Statistical Analysis

PRS were standardized with mean = 0 and SD = 1. Logistic regression was used to test for association of each psychiatric risk score with the primary phecode. The analysis was performed in R(30) with PRS as the independent variable and phecode as the dependent variable, and age, sex, and the first 10 ancestry-specific PCs as covariates. We performed logistic regression to estimate the odds ratio for cases when comparing the top quintile of polygenic risk to the remaining quintiles of risk.

PheWAS were performed for the psychiatric PRS. Logistic regression models with each PRS as the independent variable, phecodes as the dependent variables, and age, sex and the first 10 PCs as covariates, were used to identify secondary phenotypic associations. Phecodes with >100 cases (n=512) were tested. A Bonferroni-corrected phenome-wide significance threshold of p<9.77×10^−5^ was applied to account for multiple testing. To identify whether secondary trait associations of genetic risk were due to comorbid diagnoses of multiple phenotypes, we performed a conditional PheWAS, adjusting for the primary phenotype by including it as a covariate in the regression model.

## Results

### Psychiatric phenotypes in the Penn Medicine Biobank

Of the 10,182 European ancestry individuals (men n=6,676, women n=3,506, mean age = 70.4) with genotype and phenotype data in the PMBB, 3,008 (29.5%) have at least one phecode for a psychiatric disorder. Rates of psychiatric disorders are similar between men and women with 31.3% of women in the biobank having a psychiatric disorder diagnosis (n=1,097), compared to 28.6% of men (n=1,911). Table 1 details the number of cases for each psychiatric disorder, grouped by parent phecode. Tobacco use disorder is the most prevalent psychiatric disorder phecode, followed by anxiety disorders and mood disorders.

**Table 1.** Demographics of psychiatric disorders as defined by parent phecode in the Penn Medicine Biobank.

| Phecode | Description | # individuals (%) | Age |  | % Female |
| --- | --- | --- | --- | --- | --- |
|  |  |  | Mean | SD |  |
| 318 | Tobacco use disorder | 1,199 (11.8) | 69.4 | 11.1 | 29.6 |
| 300 | Anxiety, phobic and dissociative disorders | 1,121 (11.0) | 65.6 | 13.7 | 47.2 |
| 296 | Mood disorders | 1,095 (10.8) | 66.8 | 13.0 | 45.2 |
| 292 | Neurological disorders | 462 (4.5) | 73.9 | 12.0 | 31.2 |
| 317 | Alcohol-related disorders | 226 (2.2) | 66.9 | 10.2 | 16.8 |
| 316 | Substance addiction and disorders | 158 (1.6) | 62.9 | 12.6 | 29.1 |
| 290 | Delirium dementia and amnestic and other cognitive disorders | 129 (1.3) | 80.7 | 8.9 | 26.4 |
| 304 | Adjustment reaction | 90 (0.9) | 66.2 | 15.5 | 47.8 |
| 291 | Other specified nonpsychotic and/or transient mental disorders | 73 (0.7) | 70.8 | 12.3 | 42.5 |
| 313 | Pervasive developmental disorders | 70 (0.7) | 58.8 | 13.6 | 34.3 |
| 293 | Symptoms involving head and neck | 51 (0.5) | 70.2 | 12.2 | 33.3 |
| 303 | Psychogenic and somatoform disorders | 42 (0.4) | 61.9 | 14.5 | 42.9 |
| 302 | Sexual and gender identity disorders | 40 (0.4) | 67.1 | 11.6 | 10.0 |
| 301 | Personality disorders | 35 (0.3) | 68.1 | 14.6 | 40.0 |
| 295 | Schizophrenia and other psychotic disorders | 33 (0.3) | 67.2 | 13.9 | 27.3 |
| 306 | Other mental disorder | 32 (0.3) | 65.0 | 13.0 | 59.4 |
| 297 | Suicidal ideation or attempt | 18 (0.2) | 61.2 | 14.5 | 38.9 |
| 305.2 | Eating disorders | 13 (0.1) | 52.3 | 18.8 | 53.8 |
| 315 | Developmental delays and disorders | 8 (0.1) | 77.3 | 10.2 | 37.5 |
| 312 | Conduct disorders | 4 (0.03) | 69.2 | 5.6 | 25.0 |
| - | Any psychiatric disorder | 3,008 (29.5) | 68.8 | 12.8 | 36.5 |

### Primary associations of psychiatric PRS

We generated PRSs for common psychiatric disorders based on publicly available summary statistics from genome-wide association studies conducted by the Psychiatric Genomics Consortium(31) (PGC, See Supplemental Table 1 for individual references). We calculated six psychiatric PRSs, including schizophrenia (SCZ), bipolar disorder (BD), major depressive disorder (MDD), cross disorder (CROSS), attention-deficit/hyperactivity disorder (ADHD), and anorexia nervosa (AN) and tested their association with their respective primary phenotypes (Supplemental Table 3). PRS_SCZ_ was significantly associated with schizophrenia (phecode 295, N cases=33, OR=1.46, 95% CI 1.01-2.11, p=0.046). PRS_BD_ was significantly associated with bipolar disorder (phecode 296.1, N cases=96, OR=1.58, 95% CI 1.27-1.95, p=3.9×10^−5^). PRS_MDD_ was significantly associated with major depressive disorder (phecode 296.22, N cases=446, OR=1.22, 95% CI 1.10-1.34, p=1.0×10^−4^). PRS_CROSS_ was significantly associated with a composite phenotype (N cases=578, OR=1.17, 95% CI 1.06-1.29, p=0.001) comprised of autism spectrum disorders (phecode 313.3) *or* ADHD (phecode 313.1) *or* bipolar disorder (phecode 296.1) *or* major depressive disorder (phecode 296.22) *or* schizophrenia (phecode 295). Neither PRS_ADHD_ nor PRS_AN_ were significantly associated with their primary phenotypes (ADHD: phecode 313.1, AN: phecode 305.2).

Case prevalence per PRS quintile was calculated for each phenotype (Figure 1, Supplemental Table 4). Absolute risk for patients in the top quintile of PRS_SCZ_ was 0.84% (0.37% in the remaining 80%), with a corresponding odds ratio of 2.86 (95% CI 1.38-5.97, p=0.005). For PRS_BD_ absolute risk for the top quintile was 2.36% (1.09% in the remaining 80%, OR=2.21, 95% CI 1.42-3.44, p=0.0004); for PRS_MDD_ absolute risk in the top quintile was 7.69% (5.79% in the remaining 80%, OR=1.42, 95% CI 1.14-1.77, p=0.002). The top quintiles of PRS_CROSS_, PRS_ADHD_ and PRS_AN_ showed no difference in absolute risk compared to the remaining 80%.

**Figure 1.**
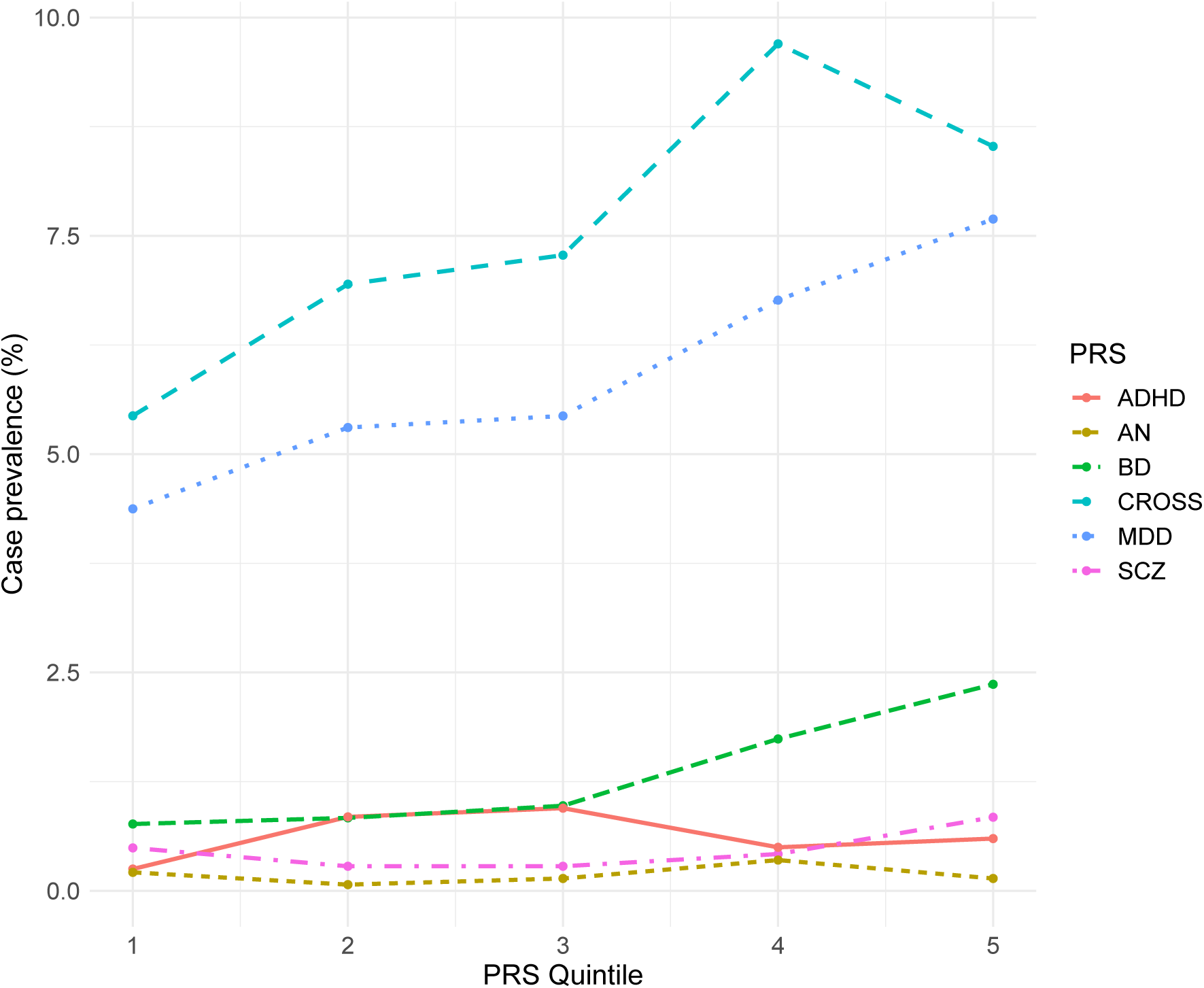
Case prevalence (%) by PRS quintile for schizophrenia (SCZ, phecode 295), bipolar disorder (BD, phecode 296.1), major depressive disorder (MDD, phecode 296.22), cross disorder (CROSS, phecodes 295, 296.1, 296.22, 313.1, 313.3), attention deficit/hyperactivity disorder (ADHD, phecode 313.1), and anorexia nervosa (AN, phecode 305.2) PRS.

### Association of polygenic risk scores with burden of psychiatric disorders

More than one-third (39.4%) of individuals (n=1,185) with a psychiatric disorder have multiple such diagnoses (Supplemental Figure 1A), as defined by the number of diagnoses for parent psychiatric phecodes within an individual. We explored the association of the psychiatric PRSs with the burden of psychiatric disorder in cases with at least one psychiatric disorder phecode. PRS_scz_ (beta=0.043, p=0.024), PRS_BD_ (beta=0.060, p=0.003), and PRS_CROSS_ (beta=0.066, p=8.98×10^−4^) were significantly associated with an increased burden of psychiatric disorder phenotypes (Supplemental Table 5, Supplemental Figure 1B).

### Phenome-wide analysis of psychiatric polygenic risk scores show cross-trait associations

To identify secondary phenotypes associated with the psychiatric PRS we conducted a PheWAS for all phecodes with >100 cases (n=512), applying a Bonferroni multiple testing correction (p<9.77×10^−5^) (Figure 2). The strongest phenotypic associations for PRS_BD,_ PRS_MDD_, PRS_CROSS_ and PRS_ADHD_ were in the psychiatric disorders domain, whereas the strongest associations for PRS_SCZ_ and PRS_AN_ were in the genitourinary and endocrine/metabolic domains, respectively. PRS_SCZ_ was significantly positively associated with “urinary system disorders” (OR=1.17, p=2.5×10^−5^) and “anxiety disorder” (OR=1.16, p=8.1×10^−5^), in addition to 11 traits at a less stringent false-discovery rate (FDR) p-value <0.05 (Supplemental Table 6). PRS_BD_ was associated with “mood disorders” (OR=1.19, p=9.3×10^−7^) and “anxiety, phobic and dissociative disorders” (OR=1.17, p=8.7×10^−6^), with associations for an additional 6 traits at FDR p<0.05 (Supplemental Table 7). PRS_MDD_ was associated with “anxiety, phobic and dissociative disorders” (OR=1.23, p=7.2×10^−10^), “mood disorders” (OR=1.21, p=1.0×10^−8^), “tobacco use disorder” (OR=1.14, p=2.5×10^−5^), and “chronic airway obstruction” (OR=1.14, p=4.4×10^−5^), as well as 8 other traits at FDR p<0.05 (Supplemental Table 8). PRS_CROSS_ was associated with “mood disorders” (OR=1.17, p=2.6×10^−5^) with an additional trait associated at FDR p<0.05 (Supplemental Table 9). PRS_ADHD_ was positively associated with “tobacco use disorder” (OR=1.27, p=8.4×10^−14^), “chronic airway obstruction” (OR=1.20, p=2.2×10^−8^) and “type 2 diabetes” (OR=1.10, p=9.5×10^−5^), and negatively associated with “benign neoplasm of skin” (OR=0.85, p=8.6×10^−9^) and “myopia” (OR=0.74, p=2.4×10^−5^), in addition to 22 traits at FDR p<0.05 (Supplemental Table 10). PRS_AN_ was negatively associated with “overweight, obesity and other hyperalimentation” (OR=0.89, p=2.7×10^−6^), and “obesity” at FDR p<0.05 (Supplemental Table 11).

**Figure 2:**
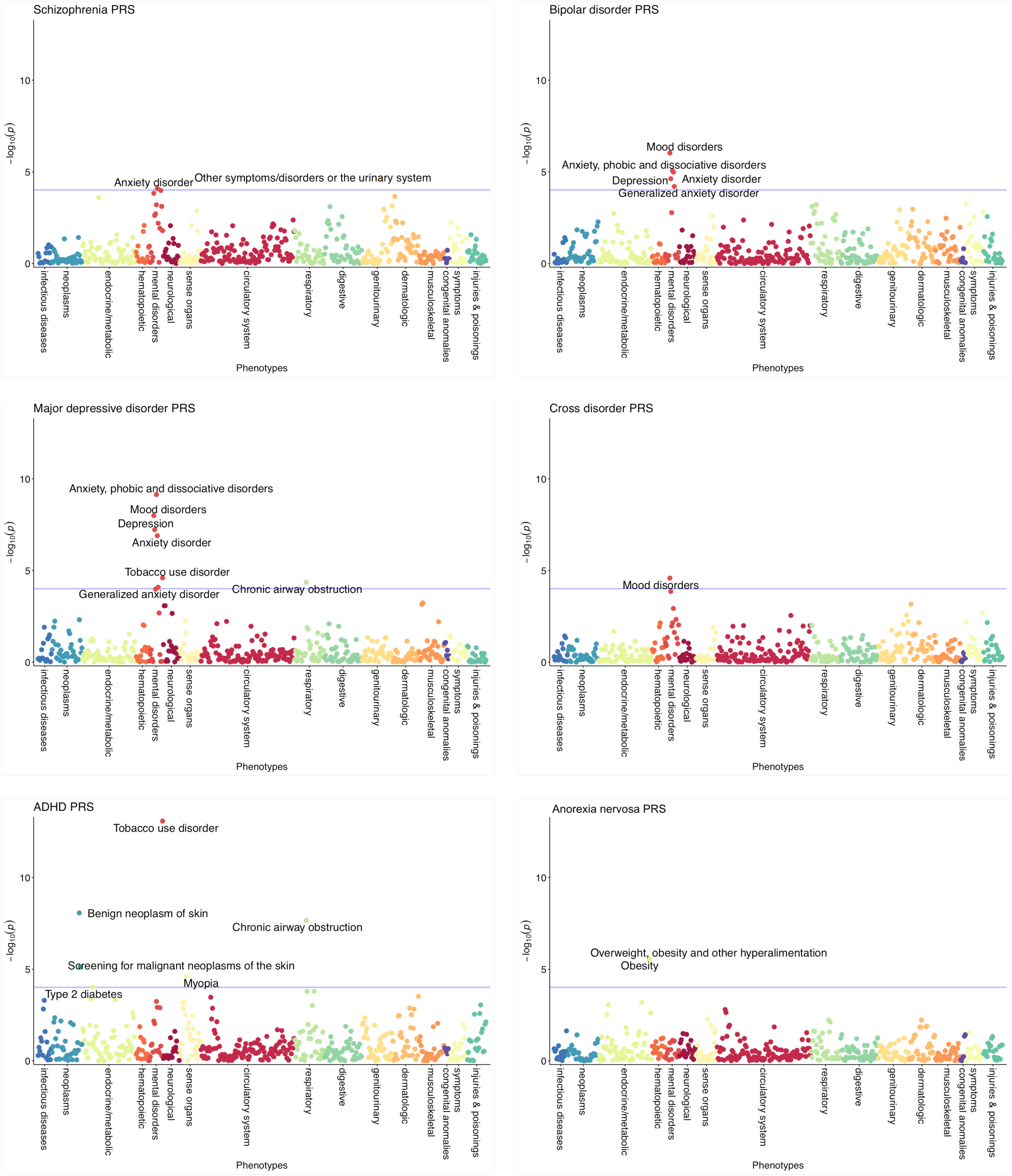
Phenome-wide association of psychiatric polygenic risk scores. A phenome-wide association plot is shown for each psychiatric PRS (schizophrenia, bipolar disorder, major depressive disorder, cross disorder, attention deficit/hyperactivity disorder, and anorexia nervosa). Phenotypes with >100 cases were tested (n=512) in 10,182 individuals from the Penn Medicine Biobank. Phenotypes are grouped by trait domain along the horizontal axis, the significance of association between the PRS and phenotype is shown on the vertical axis (-log_10_ p; two-tailed). The blue line indicates phenome-wide significance following Bonferroni correction (p<9.77×10^−5^). Phenotypes reaching phenome-wide significance are labelled.

Using an FDR threshold of p<0.05 to select significant phenotypic associations, we explored the phenotypic overlap between the psychiatric PRS (Supplemental Table 12). Almost all of the shared phenotypes were in the psychiatric disorders domain, reflecting the main associations identified above. The highest overlap was between PRS_BD_ and PRS_MDD_, with five mental disorder phenotypes (Mood disorders; Depression; Anxiety, phobic and dissociative disorders; Anxiety disorder; Generalized anxiety disorder) associated with genetic liability for both disorders. PRS_MDD_ also shared four phenotypic associations (Anxiety, phobic and dissociative disorders; Anxiety disorder; Tobacco use disorder; Chronic airway obstruction) with PRS_ADHD_. PRS_AN_ had the lowest phenotypic overlap with other psychiatric PRS. Both PRS_AN_ and PRS_ADHD_ were associated with “overweight, obesity and other hyperalimentation”, although the associations were in the opposite direction.

To identify whether secondary trait associations of genetic risk were due to comorbid diagnoses of multiple phenotypes, we re-ran the phenome-wide association analyses with the addition of a covariate for the corresponding phenotype (Supplemental Tables 6-11). The majority of phenotypic associations showed little change, with the secondary associations identified for PRS_SCZ_, PRS_BD_, PRS_ADHD_ and PRS_AN_ remaining significant. The association between PRS_MDD_ and tobacco use disorder was reduced and no longer survived Bonferroni correction, although the effect size was similar in magnitude (OR=1.13, p=1.2×10^−4^). PRS_CROSS_ was no longer significantly associated with mood disorders after Bonferroni correction (OR=1.15, p=0.003).

### PRS for other traits predict psychiatric phenotypes

Psychiatric disorders are often comorbid with other medical disorders. We therefore generated PRS for an additional 17 traits and disorders previously associated with psychiatric phenotypes, and explored their correlational structure (Figure 3A, Supplemental Table 13). PRS for psychiatric disorders were significantly positively correlated with each other, with the strongest associations being between PRS_SCZ_/PRS_CROSS_ (r=0.57), PRS_BD_/PRS_CROSS_ (r=0.53), and PRS_SCZ_/PRS_BD_ (r=0.5). The psychiatric PRSs were also positively correlated with PRS for diabetes (r=0.06 to 0.24), former smoking (r=0.08 to 0.28) and total cholesterol (r=0.0 to 0.23), and negatively correlated with birthweight (r=-0.02 to −0.11). Years of education PRS was positively correlated with PRS_SCZ_ (r=0.12), PRS_BD_ (r=0.17), PRS_AN_ (r=0.14) and PRS_CROSS_ (r=0.18), and negatively correlated with PRS_MDD_ (r=-0.09) and PRS_ADHD_ (r=-0.21). Coronary artery disease PRS was positively correlated with PRS_ADHD_ (r=0.08), and negatively correlated with PRS_CROSS_ (r=-0.11), PRS_AN_ (r=-0.04), and PRS_BD_ (r=-0.06).

**Figure 3.**
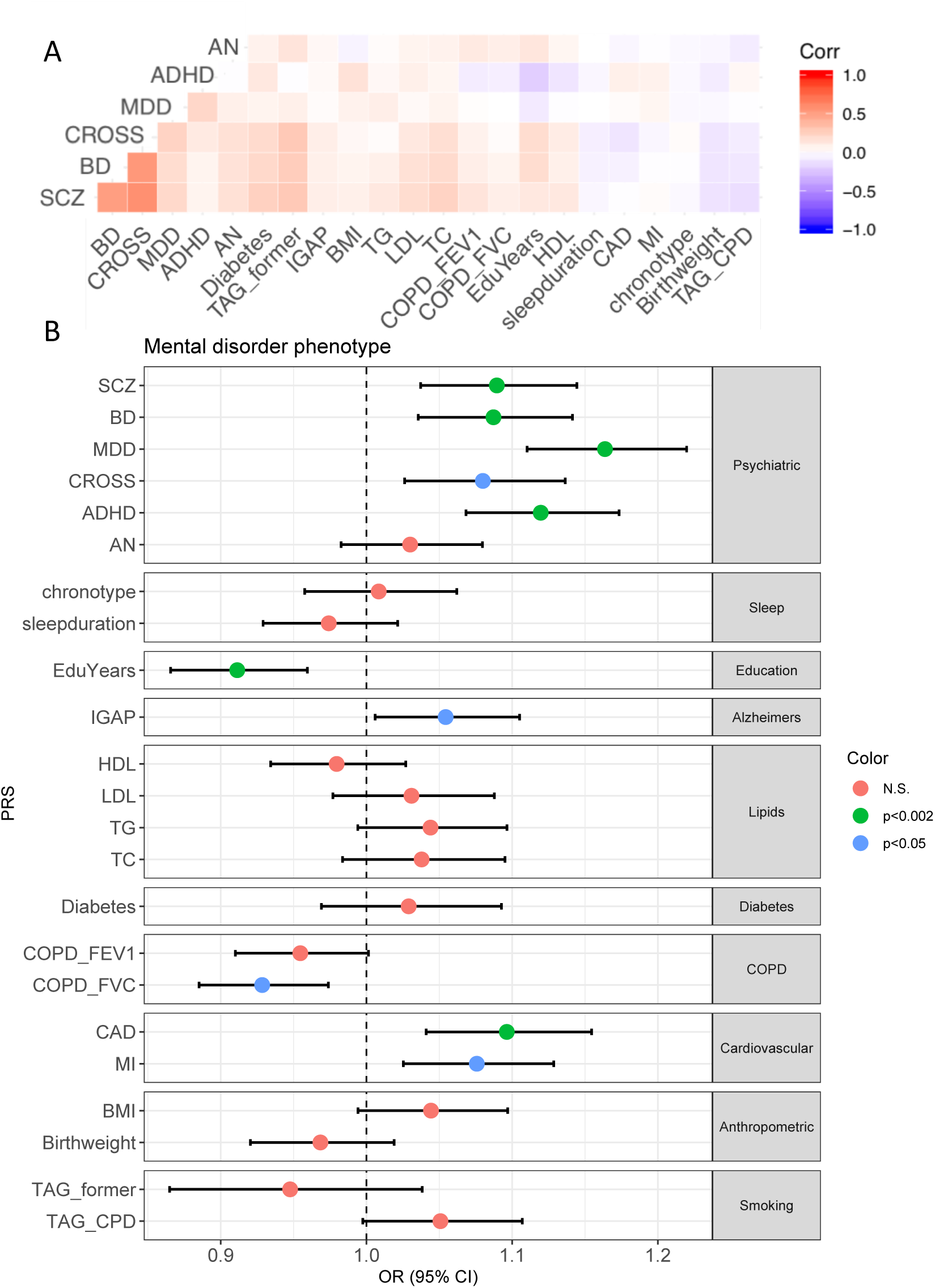
Polygenic risk for other traits. SCZ=schizophrenia, BD=bipolar disorder, MDD=major depressive disorder, CROSS=cross disorder, ADHD=attention deficit/hyperactivity disorder, AN=anorexia nervosa, chronotype=chronotype, sleepduration=sleep duration, EduYears=years of education, IGAP=Alzheimer’s, HDL=high-density lipoproteins, LDL=low-density lipoproteins, TG=triglycerides, TC=total cholesterol, Diabetes=type 2 diabetes, COPD_FEV1=chronic obstructive pulmonary disease forced expiratory volume, COPD_FVC= chronic obstructive pulmonary disease forced vital capacity, CAD=coronary artery disease, MI=myocardial infarction, BMI=body mass index, Birthweight=birthweight, TAG_former=current vs. former smoker, TAG_CPD=cigarettes per day. A: Correlational structure between psychiatric PRS and PRS for other traits. Positive correlations are denoted in red, negative correlations are denoted in blue, white squares are non-significant. B: Association between all PRS and psychiatric disorder, broadly defined. PRS are grouped by trait domain. Odds ratios and 95% confidence intervals are shown for the association between each PRS and psychiatric disorder. Psychiatric disorder was defined as an individual having any psychiatric disorder phecode.

To identify whether genetic liability for these traits was associated with a broad definition of psychiatric disorder, we identified all individuals with at least one phecode for any psychiatric disorder as defined in Table 1 (N=3,008). Of the 23 PRS tested, 6 were associated with risk for psychiatric disorder, broadly defined, at a Bonferroni-corrected p-value<0.002 (Figure 3B and Supplemental Table 14). The strongest association was with PRS_MDD_ (OR=1.16, p=2.6×10^−10^) followed by PRS_ADHD_ (OR=1.12, p=2.3×10^−6^). PRS_SCZ_ and PRS_BD_ were also significantly associated with having a psychiatric disorder (SCZ: OR=1.09, p=6.4×10^−4^; BD: OR=1.09, p=7.6×10^−4^). Two non-psychiatric PRS were associated with psychiatric disorder; PRS for coronary artery disease (CAD) was positively associated with psychiatric disorder (OR=1.10, p=5.0×10^−4^), and PRS for years of education (EduYears) was negatively associated with psychiatric disorder (OR=0.91, p=4.1×10^−4^). We conducted a secondary association test within psychiatric disorder phenotypes for PRS_CAD_ and PRS_EduYears_, and found that the most significantly associated psychiatric disorder for both PRS was “tobacco use disorder” (Supplemental Table 15).

## Discussion

Here, we present our examination of genetic liability for six psychiatric disorders with phenotypes derived from EHR diagnosis codes in 10,182 individuals from the Penn Medicine Biobank. We identify several important findings. We establish the utility of multiple psychiatric PRS in a clinical setting, showing that PRS for schizophrenia, bipolar disorder, major depressive disorder, and cross disorder are associated with their expected diagnoses in EHR. We identify cross-trait associations of psychiatric PRS mostly within the psychiatric domain, indicating overlap in the underlying genetic risk for these conditions. We identify genetic liability for two non-psychiatric disorders that increase risk of a psychiatric diagnosis.

Of the six psychiatric disorders tested, we found a significant association of genetic liability for schizophrenia, bipolar disorder, major depressive disorder, and cross disorder with their primary phenotypes, but not for attention-deficit/hyperactivity disorder and anorexia nervosa. The number of cases of ADHD and eating disorders (ED) is low in the PMBB (ADHD n=63, ED n=13), providing limited power to find a significant association. However, despite the small number of schizophrenia cases (n=33), we were able to identify a significant association between PRS_SCZ_ and schizophrenia diagnosis. It is worth noting that the schizophrenia GWAS (n=36,989 cases) is nearly twice the sample size of the ADHD GWAS (n=20,183 cases), and ten times the sample size of the anorexia nervosa GWAS (n=3,495 cases). Misclassification or under diagnosis of ADHD and ED could also be responsible for our lack of association. Cross-trait analyses for PRS_ADHD_ identified an association with tobacco use disorder, which is known to be highly prevalent in individuals with ADHD(32), suggesting that we are capturing some of the genetic risk for the disorder. Likewise, cross-trait analyses for PRS_AN_ showed a negative association with obesity, which aligns with a recent study where the authors suggest the increased liability to AN in females may be due to sex-specific anthropometric and metabolic genetic factors, specifically body fat percentage(33). In our study it is unclear as whether individuals with high PRS_AN_ have a lower propensity for obesity in the absence of an eating disorder, or whether the result is confounded by the presence of an undiagnosed eating disorder. However, this finding also suggests that we are capturing at least some of the true genetic liability for disorder.

The effect sizes of association with primary phenotypes when comparing the top quintile of psychiatric PRS to the remaining 80% (odds ratios between 1.42-2.86) are in line with previous studies of PRS in clinical settings, where non-psychiatric phenotypes including cardiovascular disease, atrial fibrillation, and type 2 diabetes have shown effect sizes in the range of 2.33-2.55(11). The schizophrenia PRS had an effect size of 2.3 in a previous study(17), compared to an effect size of 2.86 here, although our confidence intervals are larger due to the smaller sample size. Despite these medium effect sizes, the significant association of psychiatric PRS with their primary phenotypes does not provide the ability to predict the development of disorder in a naïve patient. For instance, an odds ratio for absolute risk of bipolar disorder in the top quintile of PRS_BD_ vs. the other 80% of 2.2 translates to a difference in disorder prevalence of only 1.27%. Psychiatric PRS will have to explain much more phenotypic variance and differentiate effectively cases from controls before they will be considered clinically useful. We expect that results from future GWAS will continue to improve our estimates of genetic risk, such that we can begin to consider the use of PRS as a clinical risk predictor.

PheWAS of psychiatric PRS revealed cross-trait associations, many of which were within the psychiatric disorder domain. PRS_CROSS_, representing genetic liability for five psychiatric disorders, was associated with mood disorder in a clinical setting. PRS_SCZ_, PRS_BD_, PRS_MDD,_ and PRS_ADHD_ were all associated with additional psychiatric phenotypes, suggesting a broader effect of genetic liability for these disorders. Of these, PRS_SCZ_, PRS_BD,_ and PRS_CROSS_ were also associated with an increased burden of psychiatric diagnoses, suggesting that higher genetic risk for these disorders contributes in a non-disorder specific manner to disorder load. These associations could be due to genetic liability for psychiatric disorder going beyond traditional diagnostic boundaries, or imprecision in the assignment of ICD codes at time of diagnosis. PRS_ADHD_ exhibited the most non-psychiatric phenotypic associations, including a positive association with type 2 diabetes. A positive genetic correlation between ADHD and type 2 diabetes was identified in the GWAS(7) that provided the summary statistics for the PRS; we confirm that this association can be replicated using individual level data.

Previous work showing cross-phenotype associations for PRS_SCZ_ identified many more phenotypic associations than we find here(17), likely due to the substantially larger sample size. We replicate two of these associations – anxiety disorder and urinary system disorders. Anxiety disorders were the most significant cross-phenotype association identified in Zheutlin et al. (2019), and have previously been shown to share genetic liability with schizophrenia(34). In addition, anxiety symptoms are often reported in the prodromal phase of psychoses or schizophrenia, before acute episodes, as well as being common among people with the disorder(35). We replicate the association with urinary disorders despite the fact that it was not one of the strongest associations in the prior study. Furthermore, this association remained significant in a sensitivity analysis, suggesting that it is not driven by the diagnosis of schizophrenia. Using a less stringent p-value threshold, we find many of the other phenotypic associations identified in the prior study, suggesting that these findings may be replicable across datasets.

The structure of genetic correlations we identify using individual-level PRS is largely consistent with prior genetic correlation findings for each of the psychiatric disorders. Within the psychiatric domain, PRS correlations reflect similar results to prior work showing that schizophrenia and bipolar disorder have substantial genetic overlap, and that there is moderate overlap between MDD and SCZ, BD, and ADHD(36). We also find that a genetic liability for specific disorders is associated with risk for any psychiatric disorder, defined as any diagnosis within the psychiatric disorder domain. Interestingly, we also find significant associations for genetic liability of coronary artery disease and years of education with risk for psychiatric disorder. Both of these associations appear to be driven by an association with tobacco use disorder, the largest mental disorder phenotypic group in our dataset. Despite the overall decline in tobacco use in the US over the past few decades, there are still health disparities in use and the health consequences of smoking among different social strata, including race, ethnicity, educational level, socioeconomic status, and geographic region. Notably, people with low SES are reported to smoke cigarettes more heavily, and to suffer from more diseases caused by smoking then do people from higher SES brackets. Moreover, the rate of smoking is higher among people who have a mental illness than those who do not (32.0% vs 23.2%)(37).

Our study has several limitations. The initial population available for this study contained twice as many men as women, is on average older than the general population, and a large proportion had cardiovascular disease due to the initial PMBB recruitment strategy that focused on patients with such disease (though it now recruits broadly from the health system). As researchers using EHR continue to expand and recruit more broadly from the population they serve, we expect that biobanks will become more reflective of the clinical population. Importantly, we limited our analysis to European ancestry individuals only due to the lack of GWAS summary statistics for other populations. As this area of research continues to develop, it will be critical for GWAS to be conducted in diverse ancestries to allow the clinical implications of genetic risk to be tested across populations.

We show that genetic liability for schizophrenia, bipolar disorder, major depressive disorder, and cross disorder is associated with psychiatric phenotypes derived from EHR, with effect sizes comparable to other clinical risk factors. We further conclude that genetic liability for these disorders is broadly associated with multiple psychiatric phenotypes in addition to phenotypes in other disease domains. These pleiotropic effects support evidence of an extensive shared genetic architecture among common traits. As identification of genetic markers associated with psychiatric disorders continues to improve, our results provide evidence that the application of PRS is a promising future approach for risk prediction of psychiatric disorders and associated conditions in clinical settings.

