## Supplemental Figure 1 for "Polygenic risk of psychiatric disorders exhibits cross-trait associations in electronic health record data"

### Supplemental Figures

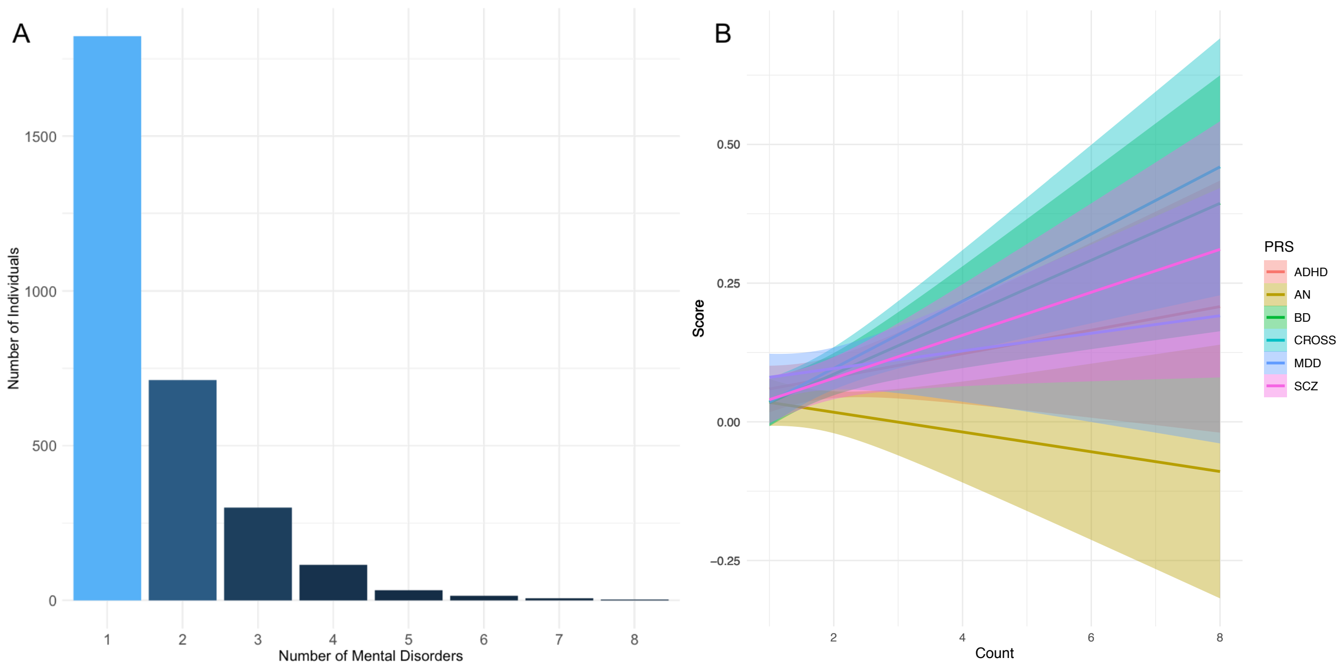


Supplemental Figure 1. A: Number of individuals with at least one parent phecode for mental disorder. B: Association of polygenic risk scores for schizophrenia (SCZ), bipolar disorder (BD), major depressive disorder (MDD), cross disorder (CROSS), ADHD (ADHD), and anorexia nervosa (AN) with burden of psychiatric disorders.
